# Identifying novel B-cell targets for chronic inflammatory autoimmune disease by screening of chemical probes in a patient-derived cell assay

**DOI:** 10.1101/2020.01.08.898270

**Authors:** Yvonne Sundström, Ming-Mei Shang, Sudeepta Kumar Panda, Caroline Grönwall, Fredrik Wermeling, Iva Gunnarsson, Ingrid E Lundberg, Michael Sundström, Per-Johan Jakobsson, Louise Berg

## Abstract

B-cell secretion of autoantibodies drives autoimmune diseases, including systemic lupus erythematosus and idiopathic inflammatory myositis. Few therapies are presently available for treatment of these patients, often resulting in unsatisfactory effects and helping only some of patients. We developed a screening assay for evaluation of novel targets suspending B-cell maturation into antibody secreting cells, which could contribute to future drug development. The assay was employed for testing 43 high quality chemical probes and compounds inhibiting under-explored protein targets, using primary cells from patients with autoimmune disease. Probes inhibiting bromodomain family proteins and histone methyl transferases demonstrated abrogation of B-cell functions to a degree comparable to a positive control, the JAK inhibitor tofacitinib. Inhibition of each target rendered a specific functional cell and potential disease modifying effect, indicating specific epigenetic protein targets as potential new intervention points for future drug discovery and development efforts.

## INTRODUCTION

B cells and autoantibodies are important drivers of autoimmune inflammatory diseases. Autoantibodies are central in the pathology of systemic lupus erythematosus (SLE) and are postulated to drive inflammation also in idiopathic inflammatory myositis (IIM)[1]. Yet, B cells are also essential in regulating immune reactions by presenting antigens to T cells and by secreting immune modulatory cytokines. SLE patients can display a range of autoantibodies to nuclear antigens including DNA, ribonucleoprotein, and chromatin proteins. IgG-autoantigen immune complexes have been hypothesized to be immune-stimulatory by activation via Fc [2] and toll like receptor (TLR) [3] pathways, and are deposited in tissues causing activation of complement. Peripheral blood B cells in SLE have been reported to be phenotypically and functionally abnormal in several ways [4, 5]; for example, there is an increased frequency of double negative memory B cells lacking CD27 in circulation [6, 7]. These IgD^-^CD27^-^ double negative memory B cells are associated with disease activity as well as with renal disease in SLE [7]. In addition, extrafollicular maturation processes of B cells have, in mouse models, been shown to have pathogenic relevance in different phases of autoimmune disease [5]. These phenotypic and functional alterations strongly suggest B cells as main drivers of autoimmune pathology in SLE.

Consequently, several B-cell targeting therapies have been evaluated and some have moved forward to clinical development [8]. B-cell depletion by the anti-CD20 targeting monoclonal rituximab is approved for treatment of patients with rheumatoid arthritis and anti-neutrophil cytoplasmic antibody (ANCA)-vasculitis and is commonly used off-lable in several autoimmune conditions, but failed to meet the efficacy end-point in SLE. In contrast, blocking B-cell activating factor BAFF (Belimumab), reducing B-cell survival, represents the only biological drug that is currently approved for treatment of SLE. Blocking BAFF mainly targets mature B cells, but other inhibitors are under clinical development for A proliferation-inducing ligand (APRIL) and its receptor that could also target long-lived plasma cells. Other strategies include monoclonal antibodies and chimeric antigen receptor (CAR)-T-cells targeting several B-cell subtypes (CD19, CD20 and CD22 expressing cells) while sparing mature long-lived antibody-secreting plasma cells. Recently, velcade/bortezomib, a small molecule proteasome inhibitor, was shown to deplete plasma cells and ameliorate refractory SLE [9], although clinical development is hampered by association with adverse events [10]. Other approaches, such as CD40 and CD40L blocking therapies, may also prove useful in regulating B-cell functions in autoimmune diseases. While these efforts are promising, there is presently only a few therapies available for the treatment of patients with SLE and IIM, often resulting in partial responses and only helping a fraction of the patients.

Hence, there is an unmet medical need in B-cell driven autoimmune diseases, including SLE and IIM. Patient derived primary cells (*e.g.* xenografts, organoids, tissue slice cultures, short term cell lines) have been explored *in vitro* for new exploratory treatment strategies for cancer, however such approaches are not commonly used in drug discovery efforts for chronic inflammatory conditions. Instead, drug development mainly in SLE but also to some degree in IIM (inclusion body myositis and polymyositis) depends primarily on studies of animal models. The clinical relevance of animal models can be questioned [11], and we believe that phenotypic screens using primary cells from patients are better suited for translational research from bed to benchside and then back to bed, for translational target evaluation.

The Structural Genomics Consortium (www.thesgc.org), a non-profit public-private organization that aim to accelerate discovery of new medicine through open research, has during the last decade developed chemical probes in collaboration with partners from the pharmaceutical industry. These small inhibitory molecules are selective, potent, and are made available to the scientific communitywithout restrictions on research use. The SGC probes target mainly epigenetic regulators, but also other proteins with critical immune functionality are contained within this collection (https://www.thesgc.org/chemical-probes).

I this study, our aim was to screen these chemical probes in a patient-derived primary B-cell assay for functions relevant in autoimmune diseases (SLE and IIM). Thus, we developed an assay using peripheral blood mononuclear cells (PBMCs) from patients and healthy donors and investigated the effect of 43 probes and compounds on induced B-cell maturation and antibody secretion. Our results show that nine of these novel probes, which inhibit bromodomain family proteins and histone methyl transferases, demonstrate abrogation of B-cell functions to a degree comparable to a positive control, the JAK inhibitor tofacitinib. We also discuss small-molecule targeting of epigenetic regulators as a potiential B-cell modulating therapeutic strategy in systemic autoimmune disease.

## MATERIAL AND METHODS

### Patients and healthy subjects

Peripheral blood was collected from 20 blood donors including five patients diagnosed with SLE and fulfilling the 1982 revised ACR classification criteria for SLE [12] and/or the SLICC-2012 criteria [13] (SLE, age 36-73 years, all females, disease duration 4-36 years), eight with IIM (7 with polymyositis and one with dermatomyositis, age 36-74 years, 6 females, disease duration 5-19 years) and seven healthy donors. All patients provided informed consent and the study was approved by the Stockholm ethical review board, in line with the Code of Ethics of the World Medical Association (Declaration of Helsinki). Two of the IIM and one of the SLE patients were untreated. Ongoing treatment of others included corticosteroids (four IIM and three SLE patients), methotrexate (two SLE), hydroxychloroquine (two SLE), mycophenolic acid (two IIM), azathioprine (one IIM), belimumab (one SLE) and rituximab (one IIM). All patients attended the Rheumatology clinic at the Karolinska University Hospital, Stockholm, Sweden.

### Chemical Probes

A set of 43 chemical probes inhibiting epigenetic regulators, protein kinases and other proteins were tested in this study. The chemical probes are small molecules and met the following criteria: *In vitro* potency of <100 nM, >30-fold selectivity *vs* other subfamilies, demonstration of on-target effect in cells at 1 µM (detailed information is available at https://www.thesgc.org/chemical-probes). To validate probe effects, negative control compunds were used when available. Negative (inactive) controls have a similar chemical structure as active probes (Figure 6b) but do not bind to the primary targets.

**Figure 1.**
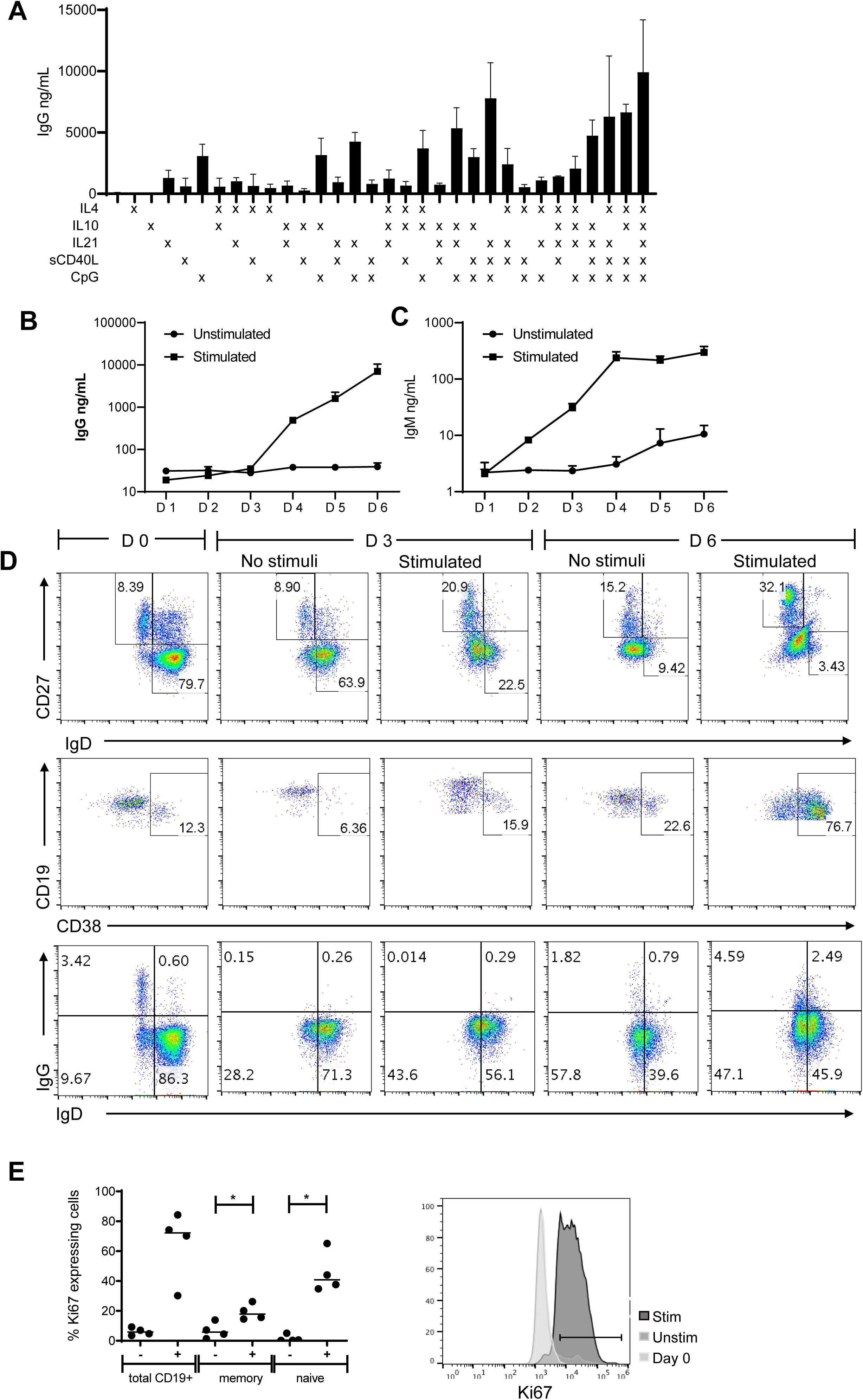
Characterization of the B-cell assay. (A) PBMCSs from healthy donors were stimulated with combinations of IL-4, IL-10, IL-21, sCD40L and CpG ODN2006 as indicated, whereafter IgG secretion was analysed by ELISA on day 6 of culture. IgG secretion (B) and IgM secretion (C) was analysed on day 1 – day 6 of culture in presence or absence of the stimulation cocktail. In A-C, average+SD of biological triplicates from one representative experiment out of three is shown. (D) B-cell phenotype was analysed on day 0, 3 and 6 of culture in absence or presence of the stimulation cocktail. Upper and lower panels show gated viable CD19^+^ B cells, while middle panels show gated CD27^+^IgD^-^ memory B cells. One experiment of two performed is shown. Numbers indicate percentage of cells in each gate. (E) Percent Ki67 expressing proliferating gated CD19^+^ B cells, CD19^+^CD27^+^ memory like and CD19^+^CD27^-^ naïve like B cells on day 6 of culture in presence (+) or absence (-) of the stimulating cocktail is shown (n=4, median is indicated, *p<0.05 Mann-Whitney). In the histogram, one example of Ki67 expression in gated CD19^+^ B cells is shown.

**Figure 2.**
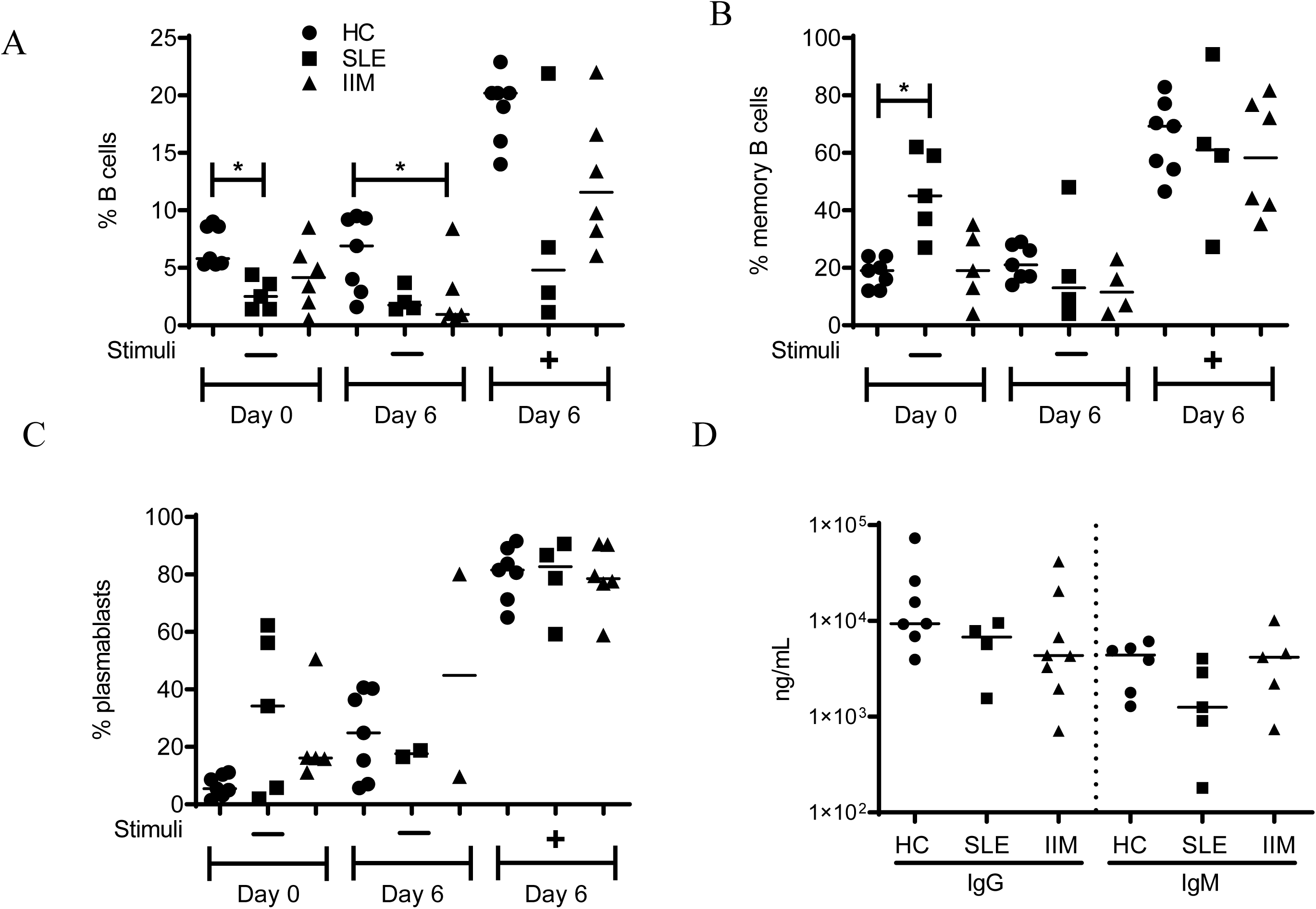
Evaluation of B-cell function and phenotype in patients with SLE and IIM. PBMCs were analyzed by flow cytometry at day 0 and day 6 of culture in absence (-) or presence (+) of the stimulating cocktail. The percentage of B cells among live CD45^+^ lymphocytes (A), the percentage of CD27^+^IgD^-^ memory cells among CD19^+^ B cells (B), the percentage of CD38^+^ plasmablasts among CD27^+^IgD^-^ memory cells (C) and the concentration of IgG and IgM in the PBMCs supernatant at day 6 of culture (D) is shown for healthy controls (HC), patients with SLE or IIM. Median values are indicated (* p<0.05 Kruskal-Wallis).

**Figure 3.**
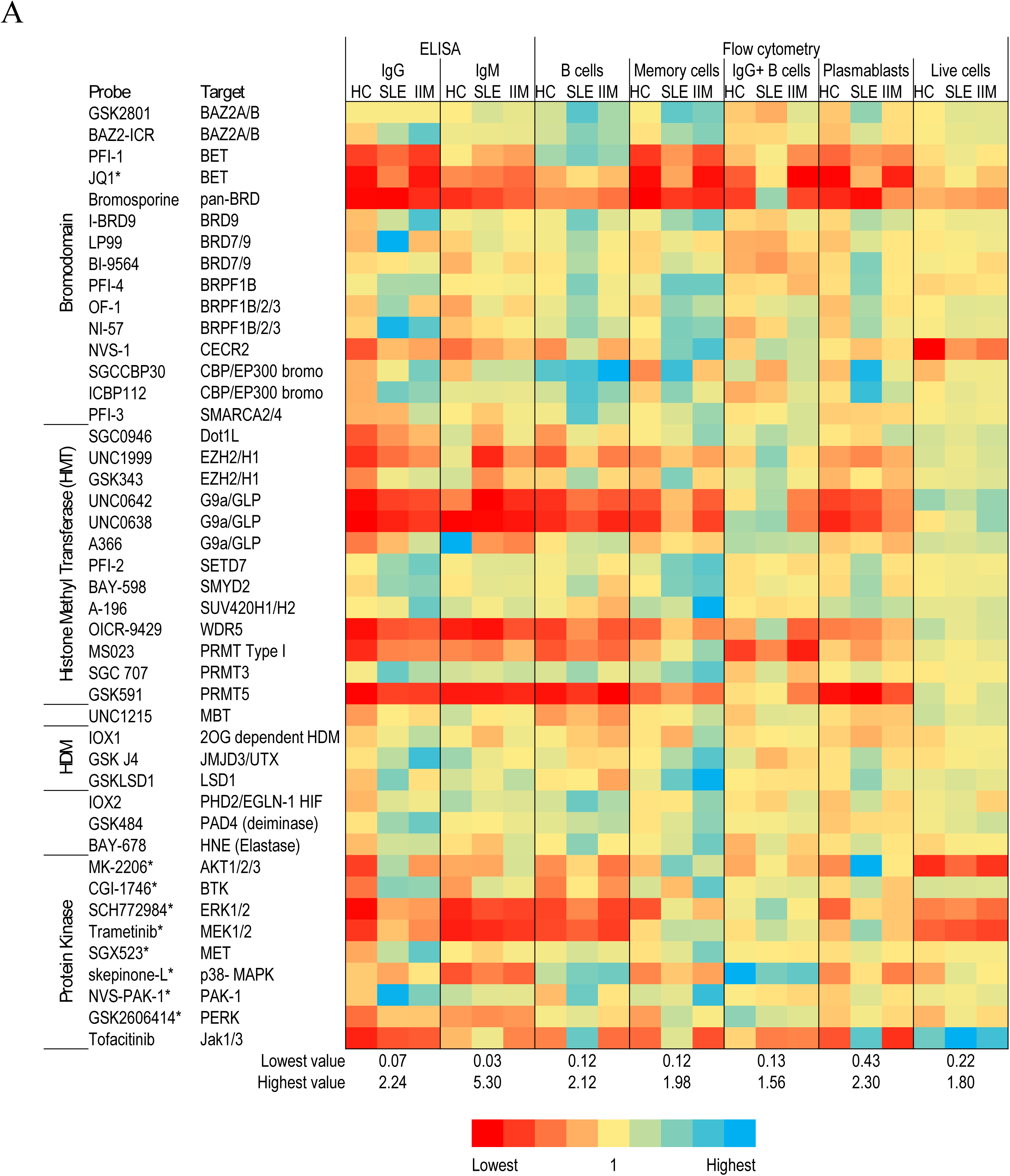
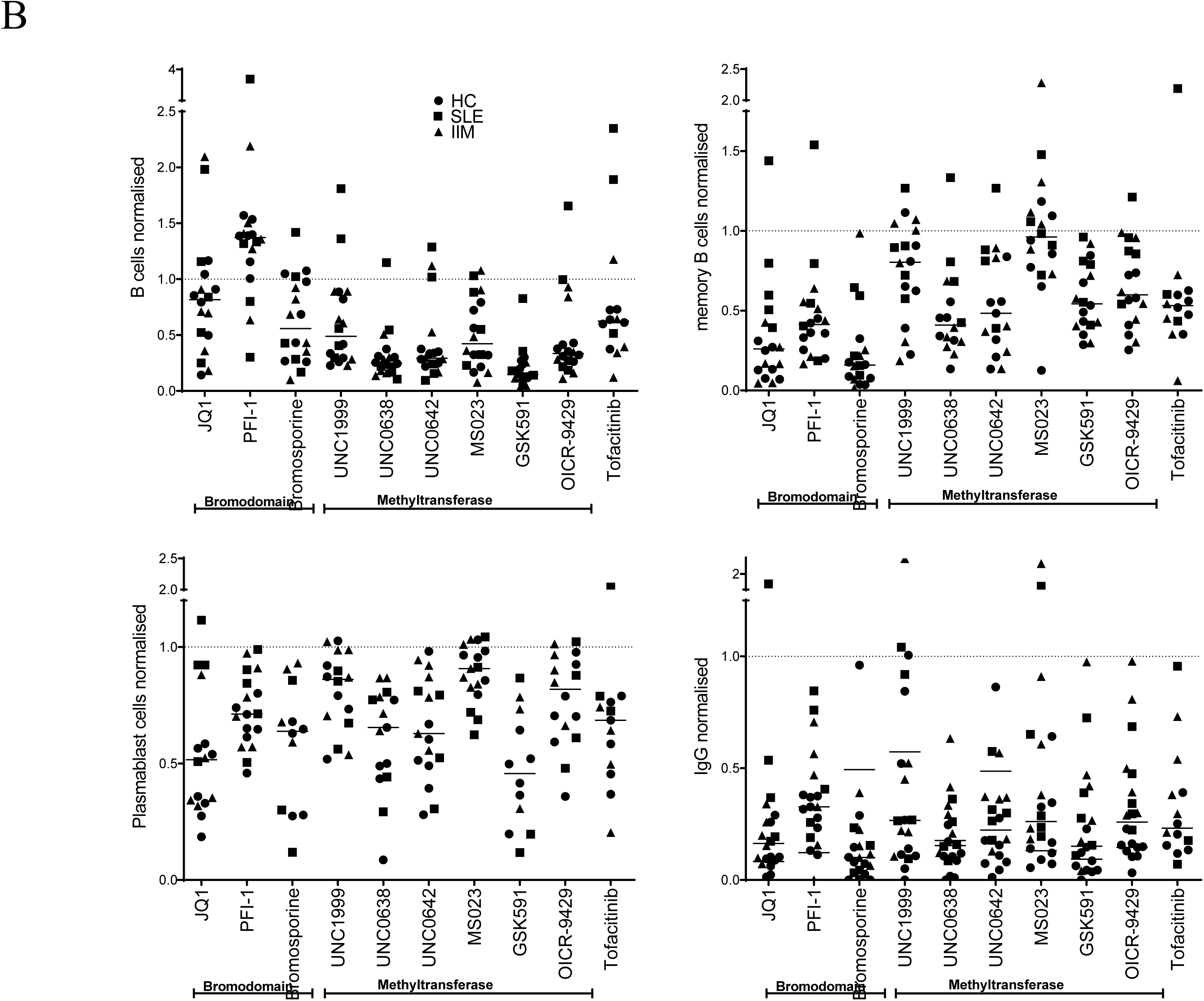
Testing chemical probes and compounds in the B-cell assay using cells from patients with SLE, IIM and healthy controls. PBMCs from healthy donors (n=7), SLE patients (n=5) and IIM patients (n=8) were cultured in presence of the stimulation cocktail, with the addition of indicated compounds for 6 days. The percentage of subpopulations of B cells and viablity of cells as analysed by flow cytometry and the secretion of IgG and IgM analysed by ELISA in presence of probes is shown normalized to DMSO vehicle. The effect of the compounds are shown by color, where blue indicates increase and red decrease of the parameter analysed. The compounds were added at 1μM (0.1μM for compounds marked with *), and target proteins are listed. In (B) individual data points of a selection of the data shown in (A) is given. Each dot is the results from one blood donor (circles healthy controls (HC), squares SLE and triangles IIM), and the median value is given. For some probes, the efficient reduction of the number of CD19^+^ B cells or CD27^+^IgD^-^ memory B cells resulted in missing data in the downstreams figures. Tofacitinib was included in 14 of the 20 experiments. Dotted lines indicate no effect compared to vehicle control.

**Figure 4.**
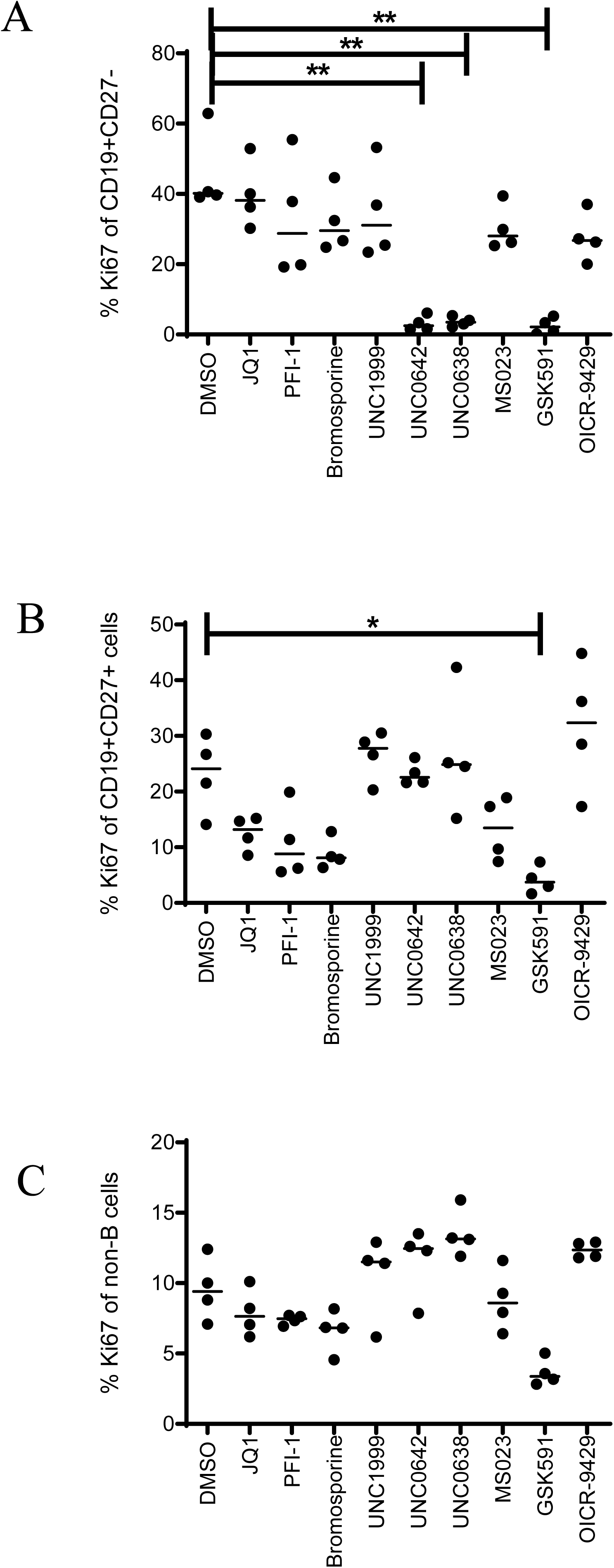
Effect of chemical probes on proliferation of B-cell subsets. PBMCs from healthy donors (n=4) were analyzed by flow cytometry for expression of the proliferation marker Ki67 at day 6 of culture in presence of the stimulating cocktail, and with the indicated compounds added at 1μM (0.1μM for JQ1). The percentage of Ki67^+^ cells among (A) CD27^-^ naïve like B cells, (B) CD27^+^ memory like B cells, and (C) CD45^+^CD19^-^ non-B cells is shown (*p<0.05, **p<0.01 Kruskal-Wallis).

**Figure 5.**
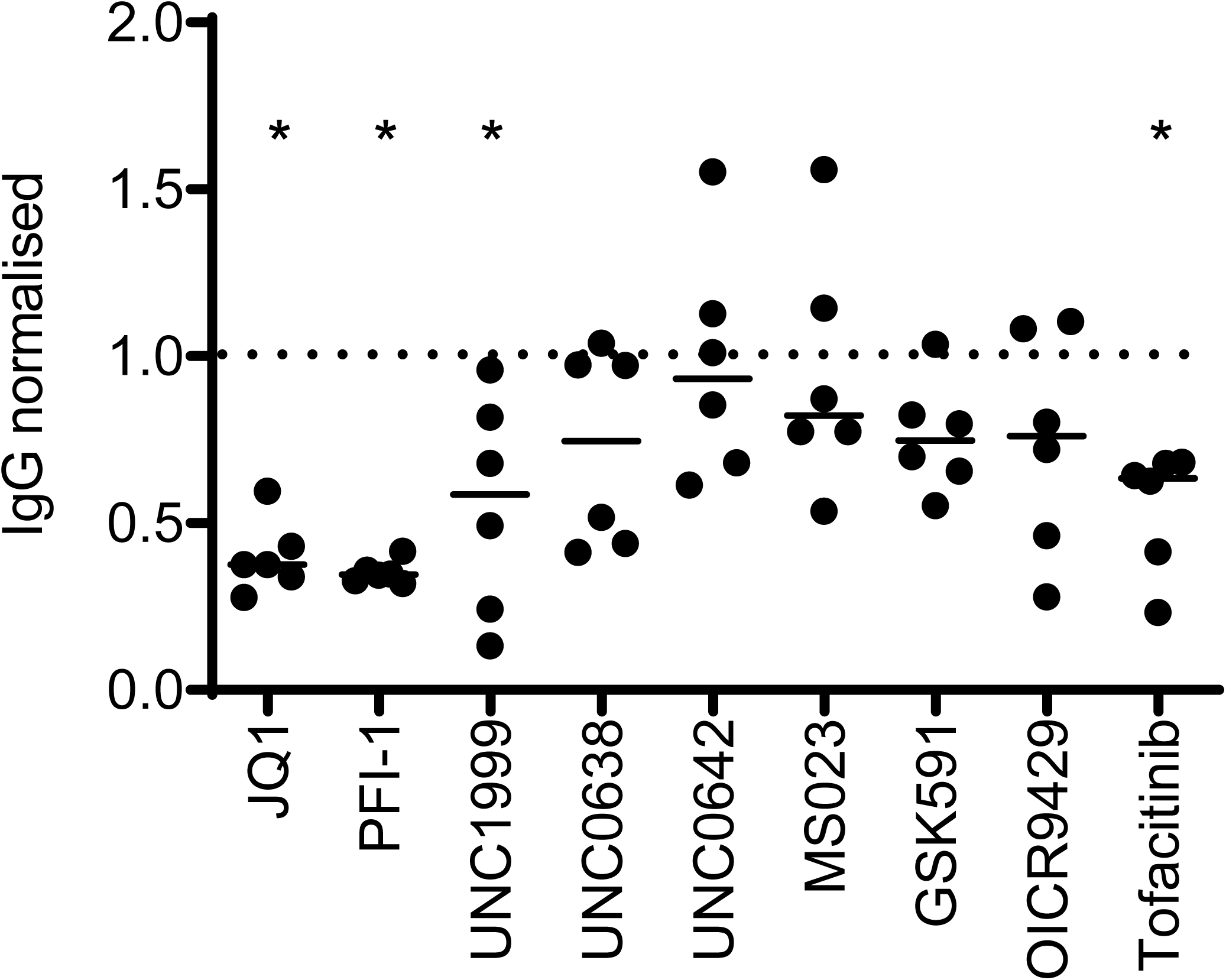
Evaluating the effect of chemical probes on *in vitro* matured B cells. PBMCs from healthy donors (n=6) cultured with the stimulatory cocktail for 6 days were washed extensively. New stimulating agents were added as well as the indicated compounds for 4 days. At the end of the 10 days, cell culture supernatants were analysed for IgG concentrations by ELISA. Median values of IgG concentration in presence of compounds are indicated as a ratio to wells stimulated in presence of vehicle control (DMSO). Dotted line indicates no effect compared to vehicle control (*p<0.05 Wilcoxon comparing absolute levels of IgG in presence of probe vs vehicle control).

**Figure 6.**
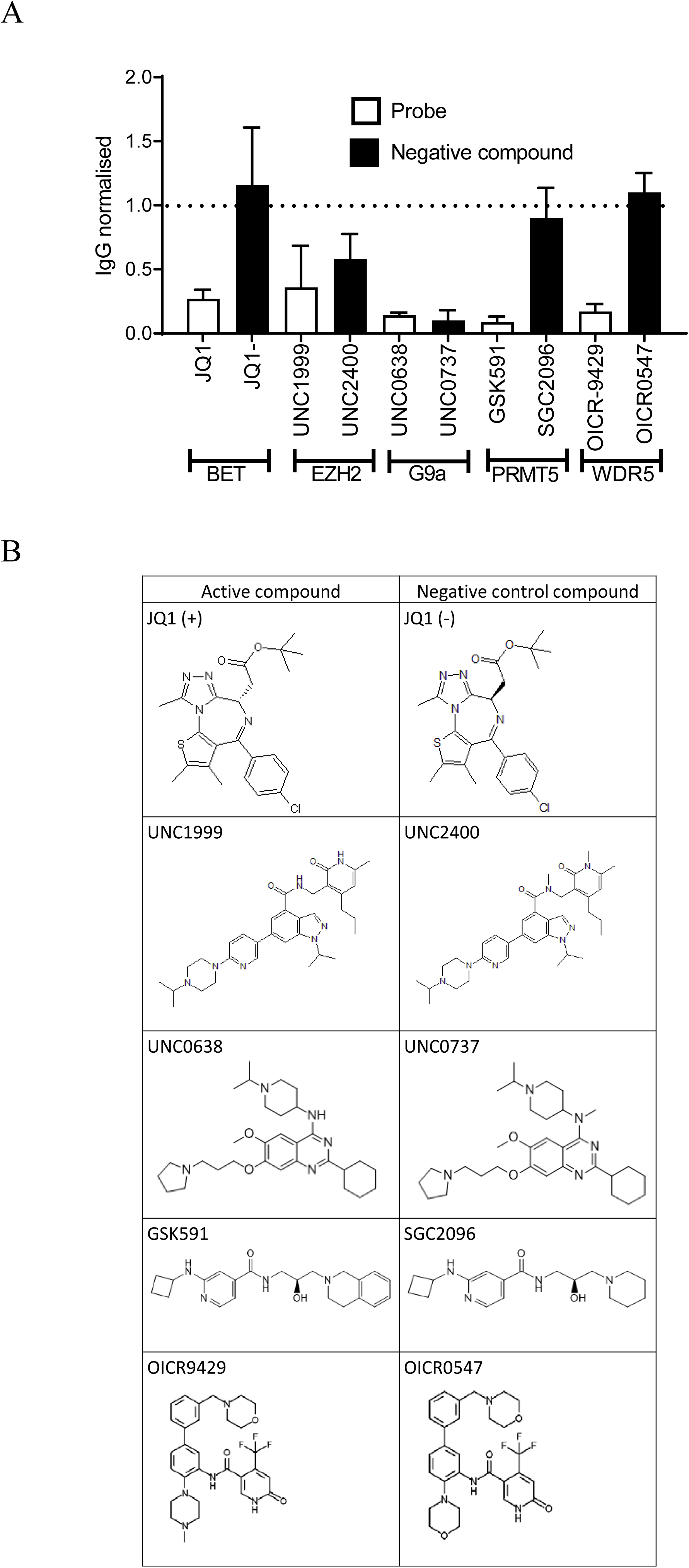
Investigating the on-target effects of probes by the use of negative control compounds. (A) PBMCs from healthy donors were cultured in presence of the stimulating cocktail and with addition of indicated compounds for 6 days. IgG secretion was analysed by ELISA, and levels of IgG in presence of probes was normalized to levels detected in presence of vehicle control (DMSO). Bars indicate the average+SD of three (two for JQ1) performed experiments. (B) Structures of active and negative control compounds.

### Isolation and in vitro culture of peripheral blood mononuclear cells (PBMCs)

PBMCs were isolated from heparinized blood or citrated buffy coat preparations by gradient centrifugation using Ficoll-Paque PLUS density gradient media (GE Healthcare Bioscience). After washing the PBMCs with the 1x Dulbecco’s Phosphate Buffered Saline (DPBS) without calcium and magnesium (Sigma Aldrich), cells were suspended in RPMI media supplemented with 10% heat-inactivated foetal calf serum (Sigma Aldrich), 1% L-glutamine (2 mM; Sigma Aldrich) and Penicillin-Streptomycin (10 ml/L; Sigma Aldrich) to a final concentration of 10^6^ cells/mL. A final volume of 200 μl/well were seeded in 96 well round-bottom plates (Sarstedt). Probes diluted in DMSO were added and plates were incubated 30 min at 37° C. Soluble rhCD40L (final concentration 500 ng/ml, R&D Systems), rhIL-21 (final concentration 50 ng/ml, R&D Systems), rhIL-10 (final concentration 200 ng/ml, R&D Systems), rhIL-4 (final concentration 200 ng/ml, R&D Systems) and CpG ODN2006 (final concentration 1 μg/ml, Miltenyi Biotec) were added, and plates were further incubated at 37° C for six days. Cells and supernatant were collected for subsequent analysis by flow cytometry and ELISA (IgG and IgM), respectively.

In a separate set of experiments, PBMCs were cultured for six days in presence or absence of stimulation cocktail as above. On day six of culture, cells were washed with DPBS twice, a new stimulatory cocktail, as well as chemical probes, were added to stimulated cells and plates incubated for a further four days. Cell culture supernatants were collected, and IgG content was determined by ELISA.

### Flow cytometry

Cells from triplicate wells were collected and pooled, washed twice with DPBS, stained with LIVE/DEAD® fixable near-IR dead cell marker (ThermoFisher), and stained for surface markers. Cells were resuspended in Stabilizing Fixative buffer (BD Biosciences) before analysis by flow cytometry. Additionally, B-cell proliferation was monitored by intracellular Ki67 AF488 (BD Biosciences) using buffers in the Fix & Perm© Cell Permeabilization Kit (BD Biosciences). Phenotypes of B cells before and after 6 days of culture were determined by flow cytometric analysis using mouse monoclonal antibody clones: CD19 PECy7 HIB19 (Biolegend), CD27 BV421 O323 (Biolegend), IgD FITC IA6-2 (BD Biosciences), Ki67 AF488 B56 (BD Biosciences), CD38 PerCPCy5.5 HIT2 (BD Biosciences), CD45 BV510 HI30 (BD Biosciences) and IgG APC IS11-3B2.2.3 (Miltenyi Biotec). Data of labelled cells were acquired using a Beckman Coulter Gallios instrument and FlowJo ™ software (BD) was used for analyzing cell populations.

### Determination of immunoglobulin concentrations

Immunoglobulins (Ig) in culture supernatants were measured using a human IgG ELISA development kit (Mabtech) according to manufacturer’s instructions, or F(ab)_2_ polyclonal goat anti-human IgM (Jackson Immuno Research) as capture antibody and detection with HRP-conjugated mouse μ-specific anti-human IgM (Southern Biotech, clone SA-DA4).

### Statistical analysis

Statistical analysis was performed using the Mann-Whitney U test or the Kruskal-Wallis test. Differences were considered statistically significant if P<0.05.

## RESULTS

### Characterization of the B-cell assay

Initially, the assay was evaluated by stimulation of PBMCs from healthy donors (n=3) with rhIL-4, rhIL-10, rhIL-21, rhsCD40L, and CpG (ODN2006) either individually or in combination, and secretion of IgG after six days of culture was assessed by ELISA (Figure 1a). Although the level of IgG produced and the level of induction using different combinations of stimuli differed between the three donors, CpG was the most important driver of IgG secretion. Hence, for the the subsequent experiments we decided to use a combination of all five stimuli. IgG production was detected above baseline from day 4 of culture (Figure 1b), while IgM induction started already at day 2 (Figure 1c).

Flow cytometry analysis of PBMCs with or without stimuli on day 0, 3 and 6 of the culture was used to investigate the effect of stimulation on B-cell phenotype. CD19-expressing B cells were identified as naïve (CD27^-^IgD^+^), memory (CD27^+^IgD^-^), IgG switched (surface IgG^+^) and plasmablasts (CD27^+^IgD^-^CD38^+^). The stimulation protocol increased the proportion of memory B cells and plasmablasts from day 3 of cell culture (Figure 1d). Expression of the proliferation marker Ki67 indicate that B cells were induced to proliferate by the stimulation. A large proportion of CD19^+^ B cells express Ki67 upon stimulation, including both memory B cells and naïve B cells (Figure 1e). B cells were rarely expressing Ki67 immediately upon PBMCS isolation as evidenced in both naive and memory B cells, (% Ki67^+^ cells n=4 median (range): 0.05% (0.03-0.3) and 0.2% (0.05-1.7), respectively). Interestingly, a substantial fraction of plasmablasts express Ki67 before culturing the cells (6.6% (0.45-18)) and the fraction of plasmablasts expressing Ki67 increases to the same level in both medium control and stimulated condition after 6 days of culture (17% (4.4-35) and 19% (7.7-22) respectively). Although the stimulation mainly induced proliferation of B cells, we noted a slight stimulation-induced Ki67 expression in non-B cells (CD45^+^CD19^-^) (2.8% (1.7-4.5)) in medium control and 9.5% (6.7-14) in stimulated cells).

### Evaluation of B-cell function and phenotype in patients with SLE and IIM

The assay was thereafter used to analyze the B-cell maturation stage and response to stimulation in PBMCs isolated from five patients with SLE, six patients with IIM andseven healthy donors. Patients with SLE showed a lower total frequency of B cells among lymphocytes in freshly isolated PBMCs as compared to healthy controls (Figure 2a), and a higher proportion of memory B cells (Figure 2b). Despite these basal differences in B-cell phenotype, stimulation induced an accumulation and maturation of B cells in both patients and healthy donors (Figure 2a-c), and IgG as well as IgM secretion was induced to a similarlevel (Figure 2d). Normalization of secreted IgG levels to the B-cell fraction of PBMCs showed similarly that the ability of the B cells to mature and produce IgG did not differ between healthy controls and patients with SLE or IMM (data not shown).

### Testing chemical probes and compounds in the B-cell assay using cells from patients with SLE, IIM and healthy controls

Next, we sought to investigate whether our defined set of chemical probes could affect B-cell phenotype and function in healthy controls and patients. We used a panel of 43 different probes and compounds at one concentration (1 μM or 0.1 μM) in the above described B-cell stimulation assay. The concentration used in our studies followed recommendations by the provider, based on experiences and results from applying these probes to various cellular assays (demonstrated on-target effect and not associated with toxicity). PBMCs from 20 blood donors including healthy donors (n=7) and patients with SLE (n=5) and IIM (n=8) were pre-incubated with probes before addition of stimuli and cultured for six days. The JAK protein kinase inhibitor tofacitinib was used as a positive control. The effect of probes was normalized to DMSO vehicle control, and an arbitrary cutoff for effect was set at (average for n=20 donors) >50% reduction of IgG secretion, without overt cytotoxic effects as analyzed by flow cytometry. Nine of the 43 chemical probes met these criteria (Figure 3) including probes inhibiting methyltransferases EZH2 (UNC1999), G9a/GLP (UNC0642 and UNC0638), WDR5 (OICR-9429), PRMT Type 1 (MS023) and PRMT5 (GSK591), as well as inhibitors of the bromodomains and extraterminal domain (BET) (PFI-1, JQ1, and Bromosporine). The results from the selected nine inhibitors and the positive control are shown for individual blood donors in Figure 3b. There was no statistically significant difference of probe effects comparing healthy donors to patient groups. However, we observed a tendency in that inhibitors of methyltransferases reduced the accumulation of B cells, whereas BET inhibitors tended to reduce their maturation into memory cells (Figure 3b). All nine probes reduced, to different degrees, the proportion of plasmablasts and IgG secretion, with similar effect as tofacitinib (Figure 3b).

### Effect of chemical probes on proliferation of B-cell subsets

Futhermore, since the stimulation cocktail induced proliferation of B cells (Figure 1e) we also investigated the effect on Ki67 expression in different B-cell subtypes. Inhibition of G9a/GLP (UNC0642 and UNC0638) and PRMT5 (GSK591) reduced the proliferation of naïve B cells (Figure 4a, p<0.01), in line with the reduced proportion of total B cells found in presence of these probes (Figure 3b). Reflecting the lower frequency of memory B cells observed following exposure to the probes (Figure 3b), BET inhibitors tended to affect the proliferation of memory B cells (Figure 4b, not significant). GSK591 clearly inhibited proliferation of total B cells compared to vehicle control (median (range) Ki67-expressing cells: 17% (1.3-32) and 73% (35-87) respectively, p<0.05), including both naïve (Figure 4a) and memory B cells (Figure 4b). None of the probes affected proliferation of CD19^-^ non-B cells significantly, while GSK591 showed a strong such tendency (Figure 4c, not significant).

### Evaluating the effect of chemical probes on *in vitro* matured B cells

Next we sought to investigate whether the probes affect the functionality of already matured B cells. Hence, we added probes to PBMCs after 6 days of stimulation, allowing B cells to mature into IgG secreting plasmablasts before inhibiting the epigenetic regulators (Figure 5). IgG secretion was induced by cytokines to a level of 4232±1711 ng/ml (Mean±SD n=6) after 6+4 days of stimulation (77±80 ng/ml without stimulation). Bromodomain targeting probes JQ1 and PFI-1 had a clear suppressive effect on IgG secretion in already matured B cells (Figure 5), suppressing by more than 60% on average. UNC1999 showed a variable suppressive effect, while tofacitinib suppressed IgG secretion by 37%.

### Investigating the on-target effects of probes by the use of negative control compounds

We then investigated whether the effects by the probes on IgG secretion in the B-cell assay were due to on-target mediated effects by including negative control compounds available for JQ1, UNC1999, UNC0638, GSK591 and OICR-9429. These compounds are similar in structure to the active compounds (Figure 6b), but do not bind and inhibit the function of the primary target protein. We were able to confirm on-target effects of GSK591, OICT-9429 and JQ1 (Figure 6a), while UNC0638 and UNC1999 displayed off-target or scaffold effects.

## DISCUSSION

Targeting B cells and their differentiation into antibody-producing plasmablasts could constitute a promising treatment strategy for autoimmune diseases [8]. In this study, we intestigated the potential of novel chemical probes targeting epigenetic regualtors and thereby identified novel druggable targets with the potential to modulate B-cell functionality also in B cells from autoimmune patients. The differentiation of mature B cells into antibody-secreting cells occurs through genetic reprogramming [14] involving epigenetic mechanisms shown to be dysregulated in autoimmunity [15]. An abnormal DNA methylation status of SLE B cells points toward the opportunity to target epigenetic regulation for novel therapeutic approaches [16, 17]. In line with this, we find that inhibition of a distinct set of epigenetic regulators reduce the ability of B cells to mature into antibody secreting cells, including those targeting BET, G9a, WDR5, EZH2 and PRMT5.

Our findings point toward epigenetic modulators as important contributors to B-cell maturation. Studies in other laboratories have shown that the transcription factor B lymphocyte-induced maturation protein 1 (Blimp1) is the master regulator of the terminal differentiation of B cells into antibody secreting cells [18-20]. The level of Blimp1 expression is regulated by the DNA binding factors Bach2 (BTB and CNC homologue 2) and Bcl6 (B-cell lymphoma 6) [21], in part by histone acetylation levels [22]. Blimp1 executes its effects by silencing target gene transcription in association with corepressors, including G9a [23] and PRMT5 [24], which we identify as potential novel targets for future B-cell therapy. In support of our findings, PRMT5, which belongs to the type II protein arginine methyltransferases, has recently been shown to affect germinal center dynamics by regulating the expression of a large number of genes [25] including Bcl6 [26]. Blimp1 is also involved in recruiting histone methylation complex polycomb repressive complex 2 (PRC2), containing the catalytic subunit enhancer or zeste homolog 2 (EZH2) [27] another hit in our screen.

In our test system, a >50% reduction of induced IgG secretion was set as the cutoff for selection of probes to be further studied, based on effects observed with the clinically used positive control, tofacitinib. Interestingly, we found that the probes with effect in our patient-derived cell assays targeted either BET bromodomain containg proteins or HMT methyl transferases. We could also observe differential effects of inhibitors targeting BET compared to HMT on B-cell maturation processes. The HMT inhibitors reduced the proportion of B cells (Figure 3b), while BET inhibitors JQ1 and PFI-1 did not. This was in part due to inhibition of B-cell proliferation, and in particular the G9a probes UNC0638 and UNC0642 as well as the PRMT5 probe GSK591 effectively blocked proliferation specifically for naïve CD27^-^ B cells (Figure 4). Similarly, G9a deficiency in mice results in reduced B-cell proliferation and lower levels of antibodies [28]. G9a functions as a methylator of histone 3, and binds Blimp1 [23] thereby blocking transcriptional repression important in the maturation process of B cells [18-20]. Contrary to the HMT inhibitors, we show that the BET inhibitors reduced the proportion of CD27^+^ memory cells (Figure 3b), and their effect on proliferation was most clearly observed in the CD27^+^ memory cell compartment (Figure 4) and on cells which had been induced to mature before addition of probes (Figure 5). The importance of BET in B-cell differentiation and antibody production has previously been observed in JQ1 treated mice which show a reduced response to vaccination [29] through ablation of Bcl6 expression.

Moreover, the WD40-repeat protein WDR5 has recently been shown to regulate proliferation of activated B cells [30], through histone methylation provided by triggering of signals through a KDM4a/KDM4C-WDR5-Cdkn2c/Cdkn3 cascade dysregulated in SLE B cells [30]. Accordingly, we saw a marked reduction in the accumulation of B cells in the presence of WDR5 inhibitor OICR9429 (Figure 3b), while the effect on proliferation as indicated by Ki67 expression was marginal (Figure 4). The transcriptional repressor EZH2 is required for maximal antibody secretion [31] and is overexpressed in SLE leukocytes [32]. While we show reduced antibody secretion in presence of the EZH2 inhibitor UNC1999 (Figure 3b), we see very little effects on proliferation (Figure 4) in contrast to other studies [33, 34]. Of note is that the negative control compound for UNC1999, UNC2400, had similar effects as UNC1999 (Figure 6) implicating off-target effects of UNC1999.

We induced B-cell differentiation to antibody-secreting plasmablasts by simultaneous stimulation of several cellular receptors, including cytokine receptors, the toll-like receptor 9 (TLR9) and the costimulatory receptor CD40. This stimulation protocol includes both T-cell dependent (IL-21, CD40L) and independent (ODN2006) stimuli. While IL-4 promotes B-cell proliferation and immunoglobulin class switch, CD40 ligation is a potent stimulator in differentiation of B cells. IL-10 promotes class switch, growth, differentiation and survival of B cells [35-37], and IL-21 is a potent T-cell derived inducer of terminal B-cell differentiation. ODN2006 belongs to class B CpG ODN agonists of TLR9, which can induce T-cell independent B-cell responses. Clearly, this rather broad stimulation protocol results in activation of several signaling pathways, including the JAK/STAT pathway for cytokines, TRAF/MAPK for CD40 ligation and MyD88/IRAK for the TLR9 agonist. In agreement with Van Belle *et al* [38], we found that *in vitro* stimulation through TLR9 was a main driver of IgG secretion in these assays (Figure 1). In summary, although this stimulation protocol represents a model system that may not reflect all fysiological features of B-cell maturation, it captures broad aspects of B-cell biology and provides a highly relevant starting point for further studies. Importantly, we developed the assay to be compatible with medium throughput (possibly high, depending on the cell source) screening of chemical probes and larger collections using patient derived primary cells. Although this pilot study included a limited number of inhibitors, we are presently developing the protocol into a 384 well plate format to be able to screen up to 200 probes per patient. This assay has therefore the potential to be used in future personalized medicine strategies for immune mediated diseases.

Since we studied B-cell differentiation in bulk PBMCs, the probe effects detected can be due to direct effects on B cells, or by effects on other cells involved in the maturation of the B cells. To investigate whether the probes directly affect the antibody secreting machinery of already matured B cells, probes were added to PBMCs pre-stimulated for six days (Figure 5). Clearly, the BET inhibiting probes JQ1 and PFI-1 affect the function of the plasmablasts induced using this experimental protocol.

Interestingly, of the five negative control compounds available, two had similar effects compared to the active compounds UNC1999 and UNC0638 (Figure 6). This indicates off-target or scaffold effects of these probes but does not preclude that the effects observed may in part be mediated through inhibition of the target protein. The targets should be further validated in B cells by inhibitors of a distinctly different chemotype and by use of alternative genetic approaches (such as siRNA and CRISPR).

Although B cells from patients with SLE and IIM are phenotypically and functionally different from healthy donors [39] (Figure 2), we saw no evident difference of the effects of probes between patients and healthy donors. This could be due to the inter patient differences in response, but also to the stimulation protocol, which induces signaling through several different intracellular pathways, leading to that more subtle differences in responses between cells from patients with autoimmune inflammation and healthy donors are masked. Our experimental platform is also useful for testing and screening of chemical probes’ effects on B cells using healthy blood, which makes it useful to research groups not directly linked to rheumatology clinics.

In conclusion, we have developed an assay for screening of novel targets in patient-derived primary cells with a focus on B-cell functions. Using this assay, we have tested the effects of 43 chemical probes and identified compounds with promising inhibitory effects. Inhibition of each target with its designated probe rendered a specific phenotypical/functional cell and potential disease modifying effect. This in turn indicates that specific epigenetic protein targets could be potential new intervention points for future drug discovery and development efforts.

## Abbreviations

ANCA: anti-neutrophil cytoplasmic antibody
APRIL: A proliferation-inducing ligand
Bach2: BTB and CNC homologue 2
BAFF: B-cell activating factor
Bcl6: B-cell lymphoma 6
BET: bromodomains and extraterminal domain
Blimp1: B lymphocyte-induced maturation protein 1
CAR: chimeric antigen receptor
DPBS: Dulbecco’s Phosphate Buffered Saline
EZH2: enhancer or zeste homolog 2
Ig: Immunoglobulins
IIM: idiopathic inflammatory myositis
PBMCs: peripheral blood mononuclear cells
PRC2: polycomb repressive complex 2
SLE: systemic lupus erythematosus
TLR: toll like receptor
WDR5: WD40-repeat protein 5

## ACKNOWLEDGEMENTS

The authors thank all the participating patients and healthy blood donors for their cooperation; all the doctors and nurses in the Rheumatology unit involved in the study; Suzanne Ackloo at SGC Toronto for support on usage and handling of chemical probes; and Fariba Foroogh and Ting Chen for technical assistance. This work has received support from the EU/EFPIA Innovative Medicines Initiative Joint Undertaking (ULTRA-DD grant no 115766). All authors have read the journal’s authorship agreement and policy on disclosure of potential conflicts of interest. The authors declare no conflicts of interest.

## AUTHOR CONTRIBUTIONS

Conceptualization: MS, LB, CG, PJJ. Methodology Development: YS, LB, CG. Formal analysis: LB, YS, MMS. Investigation: YS, MMS, LB. Resources: IL, IG, PJJ. Writing original draft: LB, SKP, YS. Writing review and editing: LB, YS, SKP, MS, CG, FW, PJJ, MMS. Visualization: YS, LB, SKP. Supervision: LB. Project Administration: LB, MS, YS. Funding acquisition: MS.

